# Regulation of transposons within medium spiny neurons enables molecular and behavioral responses to cocaine

**DOI:** 10.1101/2024.08.28.610134

**Authors:** Gabriella M. Silva, Joseph A. Picone, Amber L. Kaplan, Celeste R. Park, Diego P. Lira, R. Kijoon Kim, Natalie L. Truby, Rachel L. Neve, Xiaohong Cui, Peter J. Hamilton

## Abstract

A more complete understanding of the molecular mechanisms by which substance use is encoded in the brain could illuminate novel strategies to treat substance use disorders, including cocaine use disorder (CUD). We have previously discovered that *Zfp189,* which encodes a Krüppel-associated box zinc finger protein (KZFP) transcription factor (TF), differentially accumulates in nucleus accumbens (NAc) *Drd1+* and *Drd2+* medium spiny neurons (MSNs) over the course of cocaine exposure and is causal in producing MSN functional and behavioral changes to cocaine^1^. Here, we aimed to illuminate the brain cell-type specific molecular mechanisms through which this KZFP TF produces CUD-related brain changes, with emphasis on investigating transposable elements (TEs), since KZFPs like ZFP189 are known regulators of TEs^2–6^. First, we annotated TEs in existing single nuclei RNA-sequencing (snRNAseq) datasets of rodents that were exposed to either acute or repeated cocaine. We discovered that expression of NAc TEs was dramatically altered by cocaine experience, the most sensitive NAc cell-type was MSNs, and TEs in *Drd1+* MSNs were considerably more dynamic over the course of cocaine exposure than TEs in *Drd2+* MSNs. To determine the causality of this TE dysregulation within NAc MSNs in cocaine-induced brain changes, we virally delivered conditional synthetic ZFP189 TFs of our own design to *Drd1+* or *Drd2+* MSNs. These synthetic ZFP189 TFs are capable of directly activating (ZFP189^VPR^) or repressing (ZFP189^WT^) brain TEs^2^. We discover that behavioral and cell morphological adaptations to cocaine are produced by activating TEs with ZFP189^VPR^ in *Drd1+* MSNs or stabilizing TEs with ZFP189^WT^ in *Drd2+* MSNs, revealing a persistent opponent process balanced across MSN subtypes and weighted by TE stability and consequent gene expression within MSN subtype. We next performed snRNAseq of the whole NAc virally manipulated with ZFP189 TFs. We observed that, relative to ZFP189^WT^, NAc manipulated with ZFP189^VPR^ impeded cocaine-induced gene expression in NAc cell-types, including both *Drd1+* and *Drd2+* MSNs. Within either MSN subtype, the consequence of normal ZFP189 function was to enhance immune-related gene expression, and ZFP189^VPR^ impeded these gene expression profiles. We finally performed cocaine intravenous self-administration to determine the consequence of NAc ZFP189-mediated transcriptional control on cocaine use behaviors. We observed that ZFP189^VPR^ impeded any increases in active lever responses following a period forced cocaine abstinence. This research demonstrates that KZFP-mediated transcriptional repression of TEs within NAc MSNs is a causal molecular step in enabling gene expression and subsequent cellular and behavioral responses to cocaine use, and the use of ZFP189^VPR^ in this work demonstrates cell-type specific mechanistic strategies to block CUD-related brain adaptations, which may inform future CUD treatments.

## Introduction

Cocaine use and mortality is on the rise^7^, yet there are no FDA approved pharmacotherapies to treat cocaine use disorder (CUD). When a person uses cocaine, it enters the brain and rapidly antagonizes the dopamine transporter (DAT), which results in an increase in extracellular dopamine within the nucleus accumbens (NAc), a key brain region that encodes reward valence and drug response^8–10^. While this rapid elevation of dopamine in the NAc is associated with the euphoric effects of acute cocaine use, the long-lasting behavioral deficits associated with CUD, which exist even in the absence of cocaine use, may arise from persistent dysregulation of NAc function mediated by cocaine-induced changes in gene expression within the diverse cell-types of the NAc^11^. The principal neuronal cell-type within the NAc are GABAergic medium spiny neurons (MSNs) which can be approximately evenly divided into *Drd1+* MSNs and *Drd2+* MSNs^12,13^. The two MSN subtypes have been shown to drive distinct and often opposing responses to myriad reinforcers, including cocaine, with *Drd1+* MSN function often implicated in promoting reward and motivated behaviors, whereas *Drd2+* MSN function is often associated with decreased reward and/or aversive behaviors^14–20^. In an effort to uncover the cocaine-induced changes in gene expression that occurs within NAc MSNs, our team has previously identified that the *Zfp189* transcript differentially accumulates in MSN across the course of cocaine use and causally drives behavioral and physiological adaptations to cocaine^1^. *Zfp189* encodes the ZFP189 Krüppel-associated box (KRAB) zinc finger protein (KZFP) transcription factor (TF) of poorly understood function. *Zfp189* initially accumulates with *Drd1+* MSNs following acute cocaine exposure, and increasingly accumulates in *Drd2+* MSNs following repeated cocaine exposures, with *Zfp189* expression in either MSN subtype being sufficient to potentiate MSN electrophysiological function^1^. A separate study employing single nuclei RNA sequencing (snRNAseq) of rodent NAc demonstrated that the *Zfp189* mRNA was among the most significantly upregulated transcripts within *Drd1+* MSNs of rodents that received a single injection of cocaine^21^. Interestingly, we discovered that inducing *Zfp189* expression within NAc *Drd2+* MSNs, but not *Drd1+* MSNs, was sufficient to attenuate conditioned, reward-associated behavioral responses to cocaine^1^. This indicates that *Zfp189* accumulation within *Drd2+* MSNs may be a molecular signature of chronic cocaine exposure, which shifts the functional balance between *Drd1+* and *Drd2+* MSNs and precipitates the behavioral consequences associated with repeated cocaine use^1^. Despite these insights into the cocaine-induced time-course as well as the physiological and behavioral consequences of MSN-specific *Zfp189* expression, the *in vivo* molecular mechanisms through which this TF-encoding gene augments cocaine response is unknown.

ZFP189 is one member of the KZFP TF family, which is the largest TF gene family in vertebrates with nearly 400 distinct KZFP-encoding genes in the human genome^3^. A potential reason for this expansion of the KZFP gene family is the molecular ‘arms race’ between KZFPs and genomic transposable elements (TEs)^3,4^. TEs are self-replicating DNA sequences, many of ancient retroviral origin, that are capable of ‘jumping’ and inserting their sequence elsewhere in the genome^22,23^. KZFPs represent the major biological line of defense against this TE replication and genomic insertion, called transposition, by directly binding TEs and silencing their transcriptional activity via the KZFP conserved, transcriptionally repressive KRAB domain^4^. As a family, the KZFP class of TFs represents the major biological mechanism for controlling the expression of TEs^3,6,24,25^, including in brain^5,26,27^. Indeed, the human ortholog of ZFP189, ZNF189, has been experimentally demonstrated to bind to and regulate TEs^3^. TEs and TE-derived DNA sequences make up about half of the human genome, and there is a growing appreciation that KZFP-mediated repression of TEs has enabled the domestication of genomic TEs to the benefit of the host organism, with includes the co-option of TEs as genomic *cis-* regulatory elements, frequently as enhancers for anti-viral and immune genes, to enable complex gene expression patterns^24,28–33^.

Cocaine exposure has been shown to increase TE expression in cultured neuroblastoma cell lines^34^ and within rodent NAc^35^. In humans with CUD, genetic insertions that were the result of TE transposition were identified in medial prefrontal cortical neurons isolated from postmortem tissue from patients who died from a cocaine overdose^36^. In the context of other psychostimulants, methamphetamine-triggered TE expression and transposition was identified in the striatum and hippocampus of rodents^37^. Taken together with the observation that the expression of brain KZFPs are highly responsive to cocaine exposure^21^, there may be a molecular balancing act between substance-induced release of TEs and KZFP-mediated repression of TEs that is differentially weighted in brain cell-types over the course of cocaine use. This disruption of molecular balance between TEs and KZFPs act may causally contribute to the pathogenesis of CUD.

Here, we explored the possibility that transcriptional regulation of brain TEs is a key step in enabling neuroadaptations to cocaine use. We aimed to test the hypothesis that the function of KZFPs, like ZFP189, facilitates the pathogenesis of CUD by stabilizing genomic TEs to enable cocaine-induced gene expression patterns within NAc MSNs which causally drives behavioral and neural adaptations to cocaine. To test this, we have created synthetic ZFP189 TFs of opposing transcriptional control for conditional viral delivery to rodent NAc. We replaced the endogenous repressive KRAB moiety of wild-type ZFP189 (ZFP189^WT^) with the transcriptional activator VP64-p65-Rta (VPR) to create the synthetic TF ZFP189^VPR^. To control for the over-expression of the TF DNA-binding domain, we also removed the KRAB moiety entirely, generating a functionally inert control TF with no functional domain (NFD; ZFP189^NFD^)^2^. In our earlier published work, these ZFP189 TFs have been validated *in vitro* and in brain to demonstrate that ZFP189^WT^ is transcriptionally repressive, ZFP189^VPR^ activates transcription of ZFP189 target genes, and ZFP189^NFD^ exerts no transcriptional control at ZFP189 target genes^2^. Importantly, virally delivered ZFP189^VPR^ directly activates the expression of TEs in the rodent brain, indicating that brain TEs are directly controlled by ZFP189 *in vivo* and ZFP189^VPR^ is capable of promoting TE release in brain^2^. Here, to characterize how TEs are involved in CUD and how KZFPs participate in their transcriptional control within MSNs, we annotate TEs in existing NAc snRNAseq datasets, conditionally deliver these synthetic ZFP189 TFs to *Drd1+* and *Drd2+* MSNs, and perform snRNAseq of the NAc manipulated with these ZFP189 TFs – all in male and female rodents exposed to cocaine. We discover that transcriptional control of TEs, either by natural (ZFP189^WT^) or synthetic (ZFP189^VPR^) means, casually controls cocaine-induced gene expression in MSNs and directly drives the imbalance between NAc *Drd1+* and *Drd2+* MSNs that is a hallmark of CUD.

## Results

### Transposable elements are released in NAc cell types over the course of cocaine exposure, most prominently in *Drd1+* and *Drd2+* MSNs

To probe the degree to which cocaine exposure dysregulates the basal repression of TEs in NAc cell-types, we analyzed existing snRNAseq of rodents exposed to a single injection of cocaine^21^. From these previously published data, we annotated TEs using SoloTE^38^ and generated differentially expressed TEs (DETEs) by NAc cell-cluster in saline versus cocaine conditions (Fig. 1). We discover that, one hour following I.P. injection of 20mg/kg cocaine, many NAc cell-types experience DETE expression on a level similar or greater than DEG expression, with the *Drd1+* MSN cell-type experiencing the most dynamic TE response of all NAc cell-types with nearly all significant DETEs being dramatically up-regulated (Fig. 1a-c; DETEs in Supp. Table 1). The second most sensitive NAc cell-type, in terms of number of significantly regulated TEs, is the *Drd2+* MSN cluster, which also experiences predominantly up-regulated TEs (Fig. 1a,b,d; DETEs in Supp. Table 1). These data indicate that a single dose of cocaine rapidly releases NAc TEs, most prominently in *Drd1+* and *Drd2+* MSNs out of all NAc cell-types.

**Figure 1:**
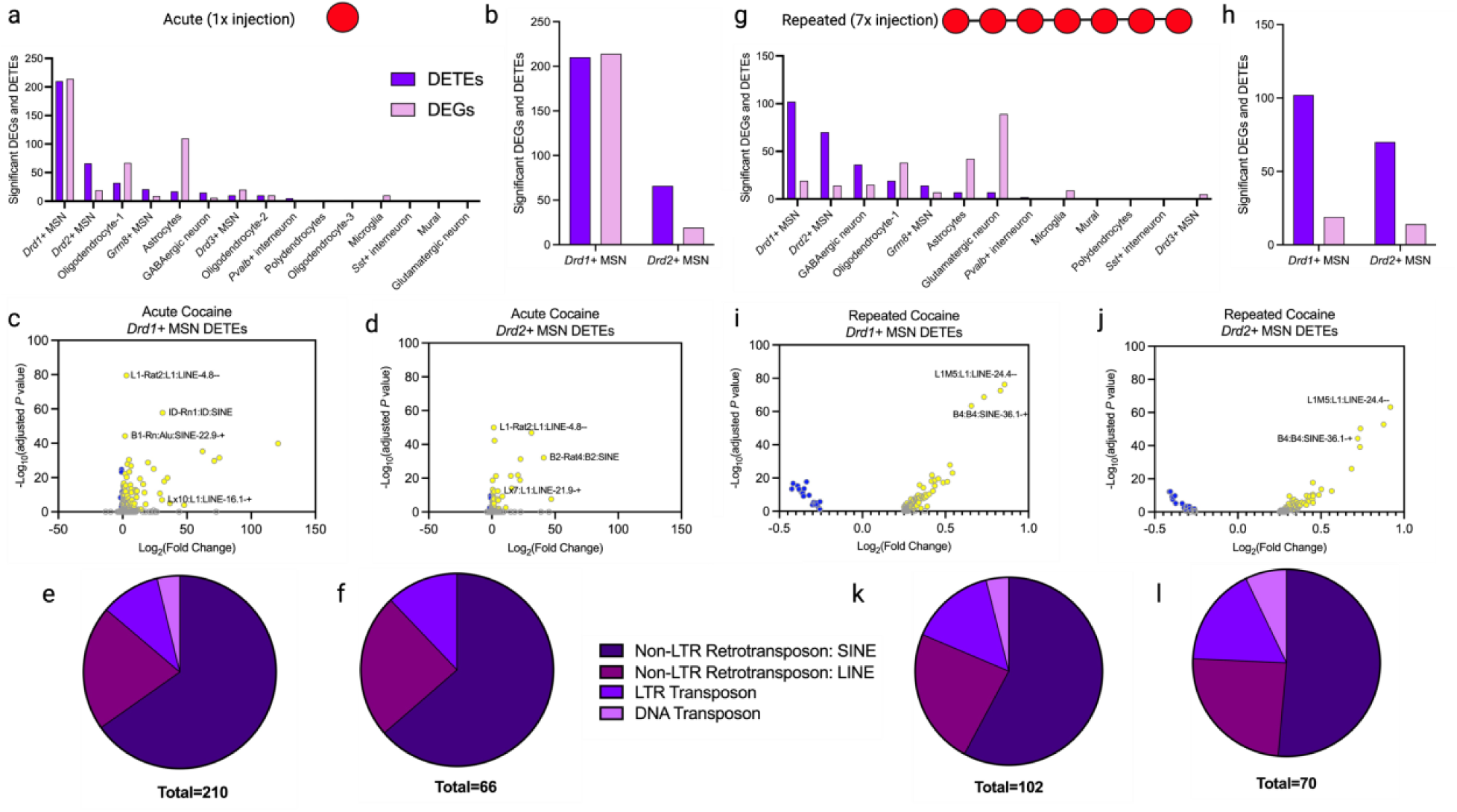
Transposable elements are released in NAc cell types over the course of cocaine exposure, most prominently in *Drd1+* and *Drd2+* MSNs. **a)** Histogram of differentially expressed genes (DEGs) and differentially expressed transposable elements (DETEs) identified within rats that received an acute dose of cocaine (single I.P. injection, 20 mg/kg). Significant DEGs and DETEs were called based on an absolute log_2_(fold change) > 0.25 with a *P*-adjusted value of < 0.05 cut-off. **b)** Histogram of MSN DEGs and DETEs. **c)** Volcano plot of DETEs within *Drd1+* MSN of rats who received an acute cocaine administration. X-axis plotted using log_2_(fold change) and Y-axis plotted using the −log_10_ (adjusted *P*-value). **d)** Volcano plot of DETEs within *Drd2+* MSN of rats that received an acute cocaine administration. **e)** Pie chart showing the amount of DETEs belonging to different transposable element families within the *Drd1+* MSN cell type. **f)** Pie chart showing the amount of DETEs belonging to different transposable element families within the *Drd2+* MSN cell type. **g)** Histogram of DEGs and DETEs identified within rats that received a repeated dose of cocaine (I.P. injection, 20 mg/kg for 7 days). Significant DEGs and DETEs were called based on an absolute log_2_(fold change) > 0.25 with a *P*-adjusted value of < 0.05 cut off. **h)** Histogram of MSN DEG and DETE regulation. **i)** Volcano plot of DETEs within *Drd1+* MSN of rats that received repeated cocaine administration. **j)** Volcano plot of DETEs within *Drd2+* MSN of rats that received repeated cocaine administration. **k)** Pie chart showing the amount of DETEs belonging to different transposable element families within the *Drd1+* MSN cell type. **l)** Pie chart showing the amount of DETEs belonging to different transposable element families within the *Drd2+* MSN cell type. This figure was generated by annotating TEs within existing and publicly available snRNAseq datasets produced by the Day Lab^21,39^.

To understand how TE regulation within NAc cell-types changes over the course of repeated cocaine exposure, we again interrogated TE expression in an additional snRNAseq study from rodents subjected to seven days of 20mg/kg cocaine I.P. injections^39^. In response to this repeated cocaine exposure, TEs are again dysregulated most prominently in *Drd1+* and *Drd2+* MSNs above all other NAc cell-types, and again mostly experience cocaine-induced increases in expression (Fig. 1g-j; DETEs in Supp. Table 2). However, the fold-change magnitude of TE up-regulation is far less, and there is an appreciable number of down-regulated TEs at this repeated cocaine time point relative to the acute time point (comparing Fig. 1c,d to Fig. 1i,j), pointing to a biological mechanism to constrain and even repress TE release that is occurring within both NAc MSN subtypes at this repeated cocaine exposure time point.

Notably, at the acute time point in all NAc cell-types, TE dynamics are positively correlated with transcriptional dynamics, meaning cell-types that possess large numbers of DETEs tend to also possess more DEGs (Supp. Fig. 1a). This suggests that dysregulation of TEs does not immediately impede gene expression. In contrast, at the repeated cocaine exposure time point, the expression of DETEs is negatively correlated with DEG expression (Supp. Fig. 1b), suggesting that the cocaine-induced activation of NAc TEs may play a role in impeding gene expression profiles within NAc cell-types at more advanced stages of cocaine exposure.

In both MSN subtypes and time points, Class I non-long terminal repeat (LTR) retrotransposons including both long interspersed nuclear elements (LINEs) and short interspersed nuclear elements (SINEs) comprise the majority, and especially the top up-regulated, DETEs detected (Fig. 1e,f,k,l; Supp. Table 1 and 2). This illustrates a common motif of LINEs and SINEs within MSN subtypes being highly sensitive to acute and repeated cocaine exposure, which is consistent with the previous literature^34–36^.

In sum, we have utilized existing snRNAseq datasets from rodent NAc to discover that Class I non-LTR TEs are rapidly and dramatically up-regulated upon a single cocaine exposure, this occurs most strongly and immediately within *Drd1+* MSNs followed by *Drd2+* MSNs; this TE release is subdued in these MSNs upon seven days of repeated cocaine exposures; and this cocaine-induced TE release may interfere with gene expression in NAc cell types upon repeated cocaine exposures. Despite these interesting findings in cocaine-induced TE dynamics within NAc MSNs, it remains unknown if the dysregulation of TEs are merely correlated with cocaine experience, or if the dysregulation of TEs plays a causative role in the pathogenic brain molecular adaptations known to accumulate upon repeated cocaine exposures. This knowledge is critical to uncover and may inform the design of pharmacotherapies to treat CUD.

### Synthetic ZFP189 transcription factors of opposing transcriptional function produce similar behavioral and molecular adaptations to cocaine when delivered to opposing MSN subtypes

To investigate how differential TE regulation within opposing MSN subtypes causally affects cocaine-evoked behavior, we conditionally introduced our novel synthetic ZFP189 TFs to the MSN subtypes of the NAc of awake and behaving mice. We achieved this by cloning the synthetic ZFP189 TFs in *lox-*stop-*lox* expression vectors, packaging in herpes simplex virus (HSV), and virally delivering to the NAc of either *Drd1-* or *Drd2-*Cre+ male and female mice via stereotaxic surgery, as we have previously to conditionally deliver other epigenetic editing constructs to targeted MSN populations in mouse NAc^1,40,41^,. Following surgery, mice were allowed to recover and then subjected to seven sequential days of 10mg/kg I.P. injection of cocaine immediately followed by a locomotor activity session in the open field arena (Fig. 2a, b). We observed that when ZFP189^VPR^ was delivered to the *Drd1+* MSNs of the NAc, animals demonstrated an immediate and persistent increase in cocaine-induced locomotion relative to other viral treatment conditions (Fig. 2c). Conversely, when ZFP189^WT^ was delivered to the *Drd2+* MSNs, animals demonstrated a similar increase in locomotion that emerged and persisted from the third day of cocaine exposure (Fig. 2d). To investigate if this change in locomotor activity was unique to cocaine exposure, we performed an identical experiment with saline injected male and female mice and observed no change in locomotion induced by any of our synthetic ZFP189 TFs (Supp. Fig. 2), suggesting that MSN-specific expression of ZFP189 TFs does not alter basal locomotion. This indicates that activation of ZFP189 gene targets within *Drd1+* MSNs, via ZFP189^VPR^, or repression of these gene targets within *Drd2+* MSNs, via ZFP189^WT^, similarly produce heightened locomotor responses to cocaine.

**Figure 2:**
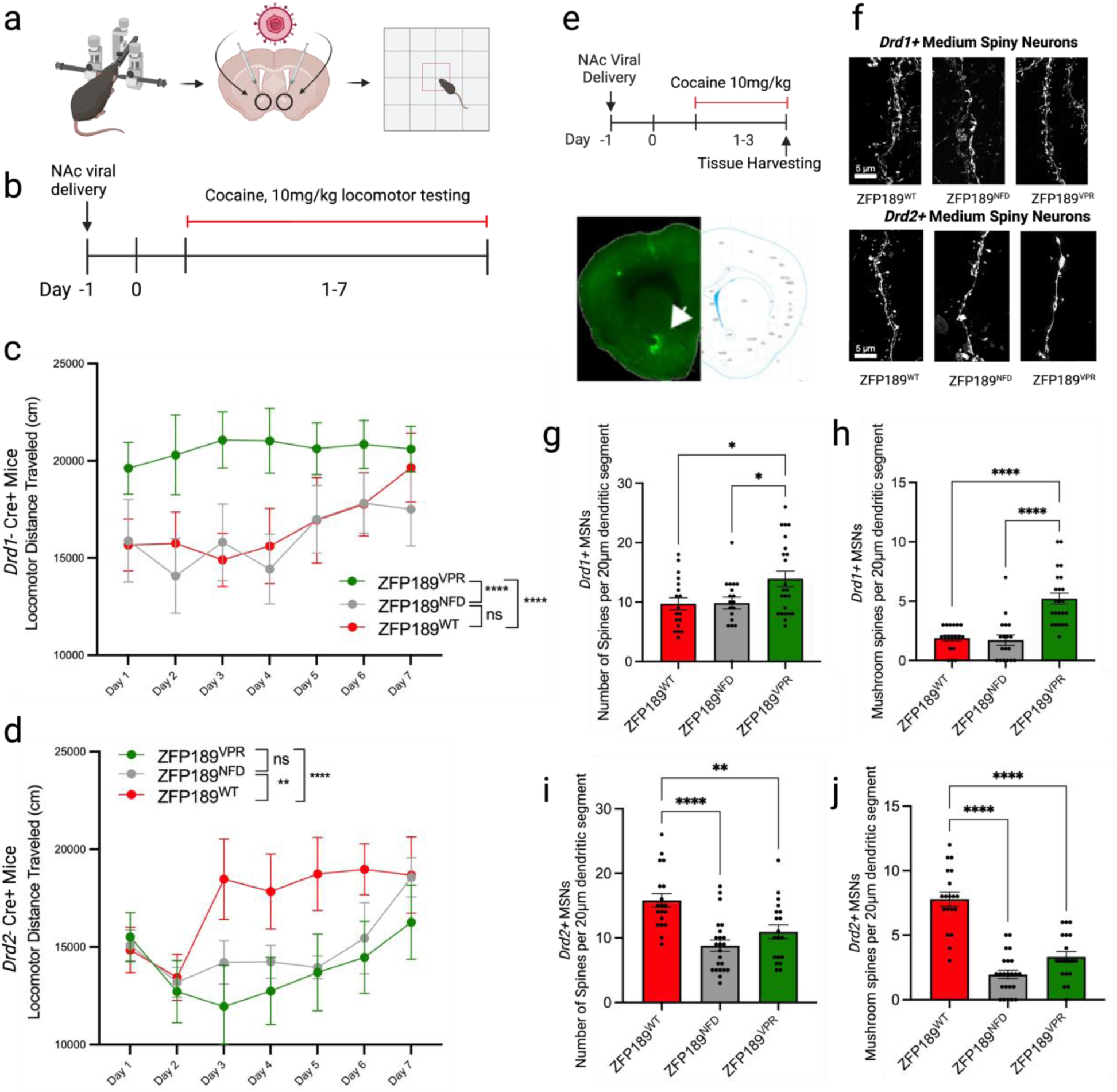
Synthetic ZFP189 transcription factors of opposing transcriptional function produce similar behavioral and molecular adaptations to cocaine when delivered to opposing MSN subtypes. **a)** Visualization of locomotor activity paradigm. **b)** Timeline of ZFP189-mediated cocaine-induced locomotor activity over a 30-minute session for seven consecutive days. **c)** Daily cocaine-induced locomotor responses within *Drd1*-Cre+ mice by ZFP189 transcription factor (TF) viral treatment, measured as distance traveled (cm). 10mg/kg cocaine, I.P. injection, 30-minute locomotor activity sessions once per day. Two-way ANOVA followed by Tukey’s multiple comparisons test, ns *P*-value > 0.05, **** *P*-value < 0.0001, n=11-12 male and female *Drd1-*Cre+ mice. **d)** Daily cocaine-induced locomotor responses within *Drd2-*Cre+ mice, by ZFP189 TF viral treatment, measured as distance traveled (cm). 10mg/kg cocaine, I.P. injection, 30-minute locomotor activity sessions once a day, Two-way ANOVA followed by Tukey’s multiple comparisons test, ns *P*-value > 0.05, ** *P*-value < 0.01, **** *P*-value < 0.0001, n=11-13 male and female *Drd2-*Cre+ mice. **e)** Timeline of ZFP189-mediated cocaine-induced spine morphology and representative targeted image. **f)** Representative *Drd1+* or *Drd2+* MSN 20μm dendritic segments from the NAc expressing one of the synthetic ZFP189 TFs and exposed to cocaine. **g)** Total number of dendritic spines counted between ZFP189 TF treatments within *Drd1+* MSNs. One-way ANOVA followed by Tukey’s multiple comparisons test, ns *P*-value > 0.05, * *P*-value < 0.05, 2-5 segments from 4-5 male and female *Drd1*-Cre+ mice. **h)** Mushroom spines counted between ZFP189 TF variants within *Drd1+* MSNs. One-way ANOVA followed by Tukey’s multiple comparisons test, ns *P*-value > 0.05, **** *P*-value < 0.0001. **i)** Total number of dendritic spines counted between ZFP189 TF variants within *Drd2+* MSNs. One-way ANOVA followed by Tukey’s multiple comparisons test, ns *P*-value > 0.05, ** *P*-value < 0.01, **** *P*-value < 0.0001, 2-5 segments from 4-6 male and female *Drd2-*Cre+ mice. **j)** Mushroom spines counted between ZFP189 TF variants within *Drd2+* MSNs. One-way ANOVA followed by Tukey’s multiple comparisons test, ns *P*-value > 0.05, **** *P*-value < 0.0001.

To further investigate the nature of how these viral manipulations affect locomotor responses to cocaine, we performed a dose response assay to probe acute behavioral sensitivity to cocaine (Supp. Fig. 3). *Drd1-* or *Drd2-*Cre+ mice were virally manipulated, exposed to a series of cumulative doses of cocaine, and locomotion was quantified. Within *Drd1-*Cre+ mice, we observed that ZFP189^VPR^ immediately potentiated sensitivity to an animal’s initial exposure to cocaine compared to ZFP189^WT^ (Supp. Fig. 3c), whereas in *Drd2-*Cre+ mice there was no change in cocaine sensitivity between ZFP189^VPR^ or ZFP189^WT^ (Supp. Fig. 3d), suggesting that dysregulating TEs within *Drd1+* MSNs, but not *Drd2+* MSNs, immediately alters an animal’s behavioral sensitivity to cocaine.

We next investigated the impact of our ZFP189 TFs on dendritic spine morphological adaptations within MSN subtypes. Spine morphology is associated with synaptic activity and spines are often classified into three morphologically distinct categories: immature thin and stubby spines versus mature stubby spines^42,43^. Upon ZFP189^VPR^ delivery to *Drd1+* MSNs in mice exposed to 10mg/kg I.P. cocaine (Fig. 2e), there was an increase in spine density and the number of mature mushroom spines on transduced *Drd1+* MSNs (Fig. 2 f-h). In contrast, when ZFP189^WT^ was delivered to *Drd2+* MSNs, there was a similar increase in spine density and number of mature mushroom spines on transduced *Drd2+* MSNs (Fig. 2 f,i,j). These manipulations had a negligible effect on the density of immature thin and stubby spines on transduced MSNs (Supp. Fig. 4), indicating that the consequence of ZFP189 molecular function within MSNs is to produce mature synaptic spines. Further, these exact viral manipulations in mice injected with saline produced MSN spine morphological consequences identical to the cocaine-treated mice in Figure 2, in that ZFP189^VPR^ delivered to *Drd1+* MSNs, and ZFP189^WT^ delivered to *Drd2+* MSNs drove the maturation of synaptic spines, even in the absence of cocaine experience (Supp. Fig. 5). Taken together, these data indicate that synthetic activation of *Drd1+* MSN TEs with ZFP189^VPR^, which induces the natural phenomenon of TE release that occurs within *Drd1+* MSNs immediately upon cocaine exposure (Fig. 1a-c), or heightened repression of *Drd2+* MSN TEs with ZFP189^WT^, which accelerates the accumulation of ZFP189 within *Drd2+* MSNs that naturally occurs upon repeated cocaine exposures^1^, both potentiate MSN structural adaptations and converge on producing heightened behavioral and dendritic spine morphological responses to cocaine. This points to KZFP-mediated transcriptional control of genomic TEs in NAc MSN subtypes is a causal molecular mechanism by which the differential contributions of these opposing MSNs are weighted to drive brain and organismal responses to addictive substances like cocaine.

### ZFP189^VPR^ impedes gene expression in diverse NAc cell-types and limits the cocaine-induced immune transcriptional adaptations in MSNs

To better understand the NAc-wide transcriptional consequences of ZFP189 function at single cell resolution, we performed snRNAseq on the NAc of mice manipulated with ZFP189 TFs and treated with cocaine. We non-conditionally delivered our three ZFP189 TFs to the NAc of male and female mice and treated them for seven days with a 10mg/kg I.P. injection of cocaine. This schedule and dose follows the exact timeline used in previous experiments (Fig. 2), and in our earlier work has been shown to cause robust transcriptional adaptations in the whole NAc by bulk RNAseq. Following the seventh cocaine treatment, mice were sacrificed, NAc were microdissected and pooled by viral treatment condition, and snRNAseq was performed (Fig. 3a). We identified 18 transcriptionally distinct clusters that could be identified as canonical NAc cell-types, including *Drd1+* and *Drd2+* MSNs, by quantifying enrichment for genes that are reported NAc cell-type markers^21^ (Fig. 3b-e). Viral treatment did not affect clustering and all cell-types were present at similar ratios, regardless of viral treatment (Fig. 3c,d). Notably, when generating DEGs within each NAc cell-type relative to the ZFP189^NFD^ treatment condition, ZFP189^VPR^ produced fewer DEGs than ZFP189^WT^, even in NAc cell-types that are not transduced with our neuron-targeting HSV delivery method, such as glial cells (Fig. 3f). In annotating TEs and generating DETEs by cell-type, we do not see ZFP189^VPR^-mediated increases in TE expression at this advanced time-point seven days post viral delivery (Supp. Fig. 7). This is consistent with the observation that TE release is transient (Fig. 1), and we have previously demonstrated that ZFP189^VPR^ activates brain TE expression 24 hours after viral delivery that could no longer be detected four-days post viral delivery^2^. Together, these data indicate that viral delivery of ZFP189^VPR^ has sweeping consequences and broadly impedes gene expression in the interconnected yet diverse cell-types of the NAc.

**Figure 3:**
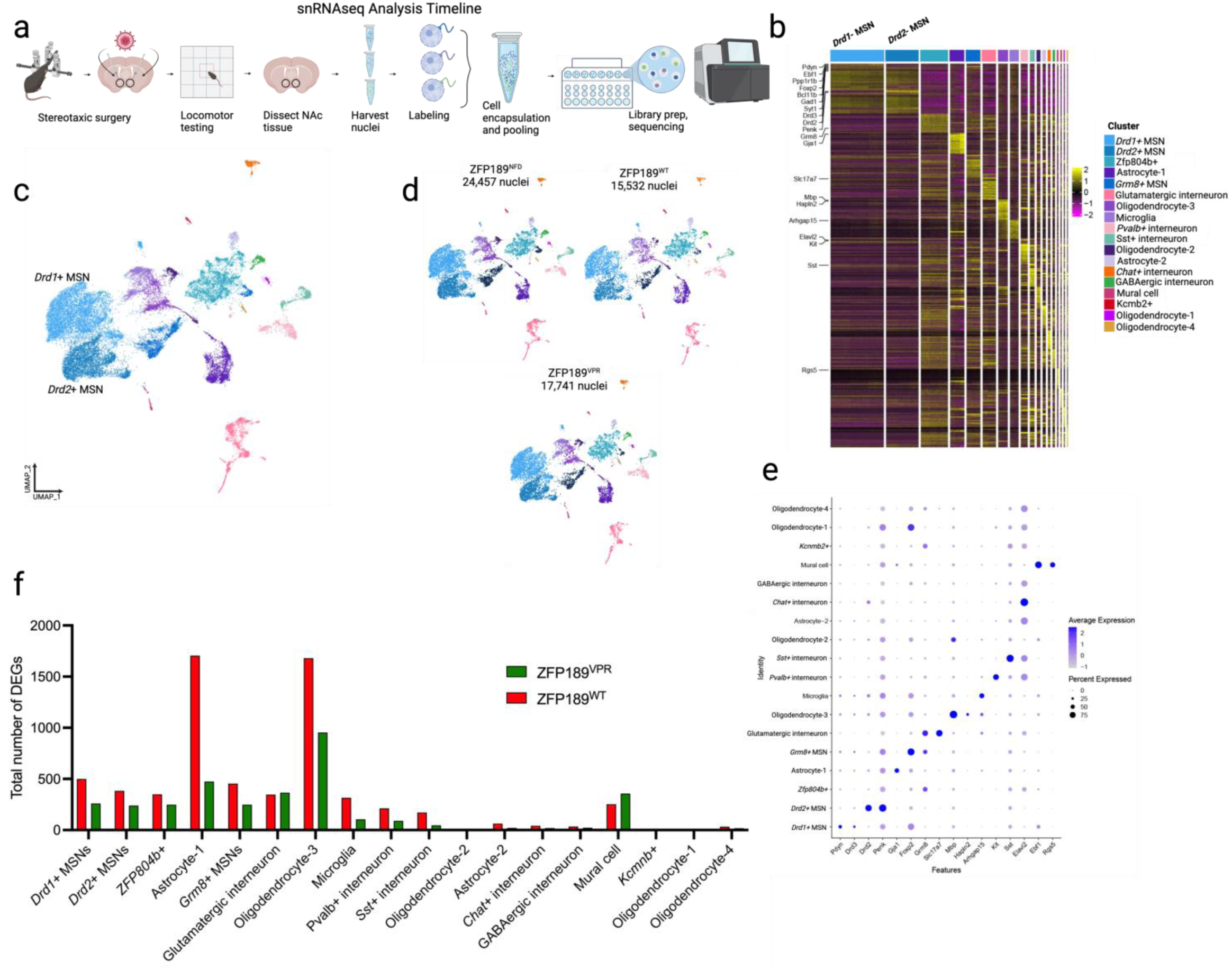
ZFP189^VPR^ impedes gene expression in diverse NAc cell-types. **a)** Timeline of single-nuclei RNA sequencing (snRNAseq) experiment. Male and female mice (n=15/sex/treatment) were delivered synthetic ZFP189 transcription factors (TFs) via stereotaxic surgery to the NAc. Mice then received seven sequential days of cocaine (10mg/kg I.P.) injection and underwent locomotor activity testing before tissue harvesting, NAc dissection, nuclei purification, and snRNAseq on the 10X Genomics platform. **b**) Heat map of cell-specific marker genes across all identified clusters. **c)** Global clustering across experimental treatment groups. NAc nuclei identifies all major classes of the mouse NAc, including *Drd1+* and *Drd2+* MSNs. UMAP, Uniform Manifold Approximation and Projection. **d)** ZFP189 TF variants similarly identify all major cell types of the NAc. **e)** Dot plot representing the average and percent expression of nuclei expressing marker genes of each identified cell-type of the NAc. **f)** Histogram of DEGs within all NAc cell types by ZFP189 TF treatment. DEGs were generated relative to the ZFP189^NFD^ control, with an absolute log_2_(fold change) > 0.25 with a *P*-adjusted value of < 0.05.

We next focused on the transcriptional consequences of ZFP189 TF function specifically within *Drd1+* and *Drd2+* MSNs. As was observed in other NAc cell-types, ZFP189^WT^ permits, and ZFP189^VPR^ impedes, the transcriptional dynamics in MSNs, as seen in the viral treatment of ZFP189^VPR^ producing far fewer DEGs in both *Drd1+* and *Drd2+* MSNs than ZFP189^WT^ (Fig. 4a). By aligning these DEGs by identity in union heat maps, we observe a similar pattern of ZFP189-regulated gene expression in both MSN subtypes (Fig. 4a). In further support of this observation, in both *Drd1+* and *Drd2+* MSNs, ZFP189^WT^ and ZFP189^VPR^ regulate largely overlapping DEGs, yet ZFP189^WT^ drives more up-regulated DEGs whereas ZFP189^VPR^ drives more down-regulated DEGs (Fig. 4b). These observations are consistent with the notion that ZFP189^WT^-mediated stabilization of genomic TEs enables *cis*-regulated gene expression, resulting in up-regulated DEGs, whereas ZFP189^VPR^-mediated activation of TEs impedes this expression of *cis-*regulated genes, resulting in fewer up-regulated DEGs of the same identity and more down-regulated DEGs (represented in the top boxed cartoon panel of Fig. 4e).

**Figure 4:**
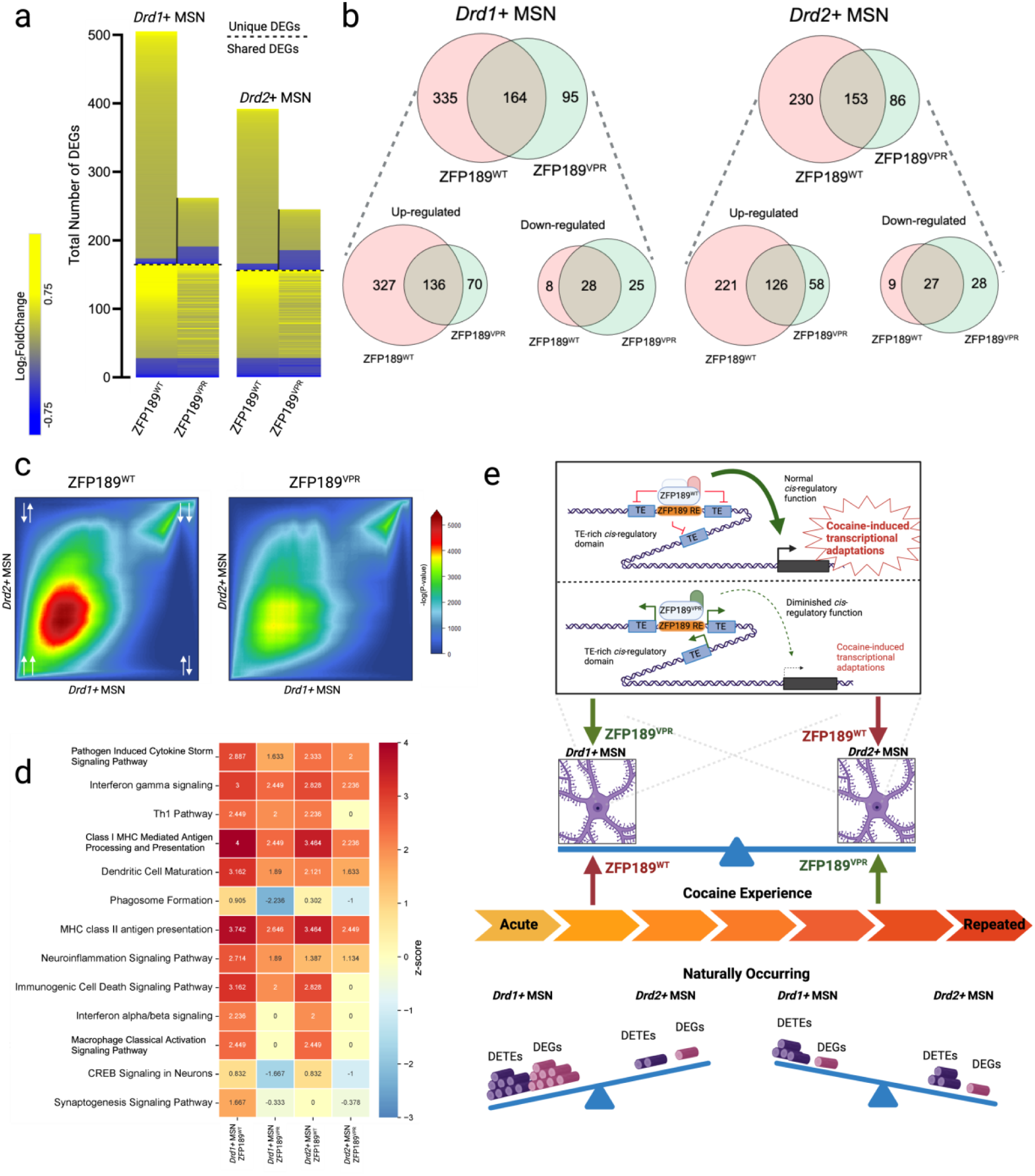
ZFP189^WT^ permits, and ZFP189^VPR^ limits, the cocaine-induced transcriptional adaptations in MSNs. **a)** Union heat maps aligned to the ZFP189^WT^ condition in MSN cell-types, showing log_2_(fold change) of genes identified as significantly upregulated (yellow) or downregulated (blue). Differentially expressed genes (DEGs) were generated against the ZFP189^NFD^ control, with an absolute log_2_(fold change) > 0.25 with a *P*-adjusted value of < 0.05. DEGs are aligned by gene ID below the dotted line, but are not aligned by gene ID above the dotted line. **b)** Venn diagrams of DEGs produced by ZFP189 TFs compared within MSN cell types. *Drd1+* MSN: chi-squared test, *P*-value < 0.0001; *Drd2+* MSN: chi-squared test, *P*-value < 0.0001. **c)** Rank-rank hypergeometric overlap (RRHO) plots mapping threshold-free DEGs produced by either ZFP189^WT^ (left panel) or ZFP189^VPR^ (right panel) within NAc *Drd1*+ or *Drd2+* MSNs. Heat indicates strength of overlap, and quadrants represent directionality of gene expression. (White arrows; bottom left: upregulated *Drd2+* MSN upregulated *Drd1+* MSN; bottom right: upregulated *Drd2+* MSN downregulated *Drd1+* MSN; top left: downregulated *Drd2+* MSN upregulated *Drd1+* MSN; top right: downregulated *Drd2+* MSN downregulated *Drd1+* MSN.) **d)** Gene ontology pathway analysis of DEGs produced by either ZFP189^WT^ or ZFP189^VPR^ within either *Drd1*+ or *Drd2*+ MSNs. Significant DEGs between ZFP189 TFs and MSN cell-types used. Significantly regulated pathways (FDR< 0.05) were sorted by z-score. **e)** Cartoon summary representing the natural dynamics of gene expression and expression of transposable elements (TEs) within either medium spiny neuron (MSN) subtype across the time-course of cocaine exposure (bottom panel). By directly manipulating the transcriptional control of TEs with synthetic ZFP189 TFs (ZFP189^WT^ or ZFP189^VPR^) in a MSN-specific manner, we manipulate a causal mechanism by which KZFP-mediated control of genomic TEs within NAc MSNs drives cocaine-induced brain adaptations.

We next investigated the threshold-free gene expression patterns induced with our synthetic ZFP189 TFs across MSN cell-types by performing a rank-rank hypergeometric overlap (RRHO). When comparing DEG profiles of ZFP189^WT^-regulated DEGs within *Drd1+* versus *Drd2+* MSNs, we see strong up-regulation of the same transcripts in either MSN subtype (Fig. 4c, left panel). Similarly, ZFP189^VPR^ up-regulates largely identical DEGs in both MSN subtypes, only to a degree less than ZFP189^WT^ (Fig. 4c, left versus right panel). This indicates that the ZFP189 TF gene targets and transcriptional consequence of ZFP189 TF expression is largely the same in either MSN subtype. Lastly, to probe the biological functions in which these affected DEGs participate, we performed Gene Ontology (GO) analyses on all significant DEGs affected by either ZFP189 TF in either MSN subtype (Fig. 4d). Within both *Drd1+* and *Drd2+* MSNs, we observe that ZFP189^WT^ drives the activation of GO terms involved in largely immune-associated pathways (Fig. 4d). Conversely, ZFP189^VPR^ affects many of these same GO terms, but to a lesser degree than ZFP189^WT^ (Fig. 4d). These results suggest that, in both MSN subtypes, ZFP189^WT^ enables the transcriptional dynamics necessary to activate the expression of predominantly immune-related transcripts. ZFP189^VPR^ affects these same genes, but restricts the transcriptional dynamics resulting in reduced activation or repression of these immune-related transcripts.

### ZFP189^VPR^ impedes cocaine self-administration behaviors

To investigate the consequence of NAc-wide transcriptional dysregulation with ZFP189^VPR^ on cocaine reward-related behaviors, we employed mouse intravenous self-administration (IVSA) for cocaine. Mice were trained to self-administer 0.5mg/kg/infusion of cocaine at a fixed ratio (FR) 1 schedule of reinforcement within three-hour daily sessions. Upon stable responding to the cocaine-paired active lever, mice were split into three viral treatment groups for delivery of one of our ZFP189 TFs. Following recovery, we recorded active lever responses for 14 daily sessions post-surgery, subjected animals to seven days of forced abstinence (FA), and then allowed the animals to return to the IVSA paradigm and resume operant responding for cocaine IV infusions for seven sequential days (Fig. 5a). In all viral treatment conditions, we observed no change in active lever responses in the sessions following viral mediated gene transfer compared to pre-surgery baseline behavior (Fig. 5c-e). However, in post-FA sessions, we saw that both ZFP189^NFD^- and ZFP189^WT^-treated mice produced a significant increase in active lever presses relative to their pre-surgery behavior (Fig. 5c-d). Interestingly, only ZFP189^VPR^-treated mice showed no change in their cocaine IVSA responses along the entire IVSA protocol, including both pre- and post-FA (Fig. 5e). Thus, by inverting the NAc molecular function of ZFP189 with ZFP189^VPR^, we block this normal behavioral escalation of cocaine taking behaviors.

**Figure 5:**
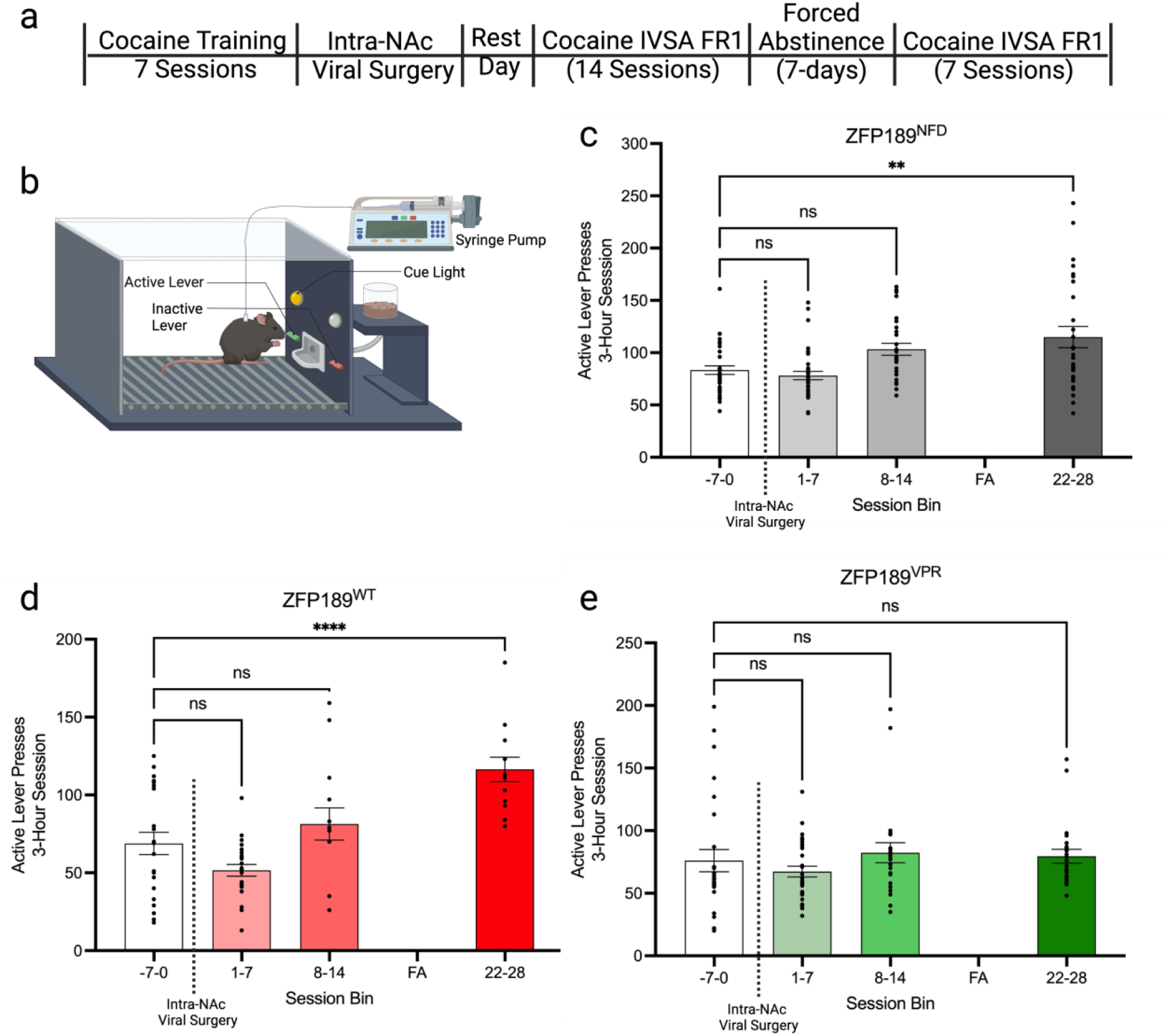
ZFP189^VPR^ impedes cocaine self-administration behaviors. **a)** Timeline of cocaine intravenous self-administration (IVSA) behavioral paradigm. Mice respond at a fixed ratio (FR) 1 for 0.5mg/kg/infusion cocaine during three-hour daily sessions. **b)** Graphic depiction of operant chamber used in IVSA. **c)** Active lever responses, grouped by seven-day session bins, for mice that received intra-NAc infusion of ZFP189^NFD^. Each dot represents an individual session. One-way ANOVA followed by Dunnett’s multiple comparisons test; ns *P*-value > 0.05, ** *P*-value < 0.01, n=5 mice. **d)** Active lever responses, grouped by seven-day session bins, for mice that received intra-NAc infusion of ZFP189^WT^. Each dot represents an individual session. One-way ANOVA followed by Dunnett’s multiple comparisons test; ns *P*-value > 0.05, **** *P*-value < 0.0001, n=7 mice. **e)** Active lever responses, grouped by seven-day session bins, for mice that received intra-NAc infusion of ZFP189^VPR^. Each dot represents an individual session. One-way ANOVA followed by Dunnett’s multiple comparisons test, ns *P*-value > 0.05 n=7 mice.

Taken together, this work points to a novel transcriptional mechanism linking KZFPs, TEs, and gene expression that similarly occurs within both NAc MSN subtypes to enable adaptations to cocaine. This molecular mechanism can be artificially manipulated by our synthetic KZFP TFs and is naturally differentially weighted across MSNs over the course of cocaine use. The consequence of this mechanism is to directly control the differential balance and functional output between the opposing *Drd1+* and *Drd2+* MSNs and produce the natural escalation of CUD-related behaviors including cocaine self-administration. This is summarized in the graphical representation of Figure 4e.

## Discussion

In this research, we discover that cocaine exposure releases TEs within NAc MSNs – a molecular phenomenon that rapidly and transiently occurs within *Drd1+* MSNs yet accumulates within *Drd2+* MSNs over the course of cocaine use. By creating synthetic ZFP189 TFs of opposing transcriptional control, we are able to intentionally either release or repress these TEs within the rodent brain^2^. By conditionally delivering these ZFP189 TFs to the *Drd1+* or *Drd2+* MSNs of mouse NAc, we discover that opposite forms of KZFP-mediated transcriptional control in these opposing cell-types converge on producing behavioral and cell morphological adaptations to cocaine. We discover that normal KZFP function is critical to produce cocaine-induced transcription within NAc MSNs, and by inverting the normal function of a KZFP with ZFP189^VPR^, we are able to impede cocaine-induced gene expression in MSNs and other NAc cell-types. This research demonstrates that KZFP-mediated transcriptional repression of TEs within NAc MSNs is a causal molecular step in enabling gene expression and subsequent cellular and behavioral responses to cocaine use.

In our previous research using CRISPR/dCas9 epigenetic editing to manipulate *Zfp189* expression in NAc MSNs^1^, it remained unclear if *Zfp189* accumulation within *Drd2+* MSNs was a singular molecular catalyst for precipitating behavioral adaptations to chronic cocaine, or if the natural *Zfp189* accumulation differentially tunes an opponent process between the *Drd1+* and *Drd2+* MSNs over the course of cocaine use. Here, by creating synthetic ZFP189 TFs of opposing transcriptional control, we are uniquely able to address this question. The fact that ZFP189^VPR^ produced behavioral and morphological adaptations when delivered to *Drd1+* MSNs but not *Drd2+* MSNs, and ZFP189^WT^ produced behavioral and morphological adaptations when delivered to *Drd2+* MSNs but not *Drd1+* MSNs, points to a persistent opponent process that is weighted by the degree to which the MSN subtype is able to control TE stability and consequent gene expression within the cell-type. This means that cocaine-induced KZFP accumulation and TE dynamics in the *Drd2+* MSNs, which we associate with chronic cocaine use, is not a molecular point-of-no-return for permanent cocaine-induced MSN changes. Rather, cocaine use shifts the weighted contributions of these opposing cell-types by differentially dysregulating and then stabilizing TEs by MSN subtype, rapidly in *Drd1+* MSNs and more steadily in *Drd2+* MSNs, over the course of cocaine use (see Figure 4e for graphical summary). Knowing this, devising therapies that could halt the cocaine-induced KZFP-mediated stabilization of TEs or invert this molecular phenomenon in targeted MSN populations, akin to what is achieved with ZFP189^VPR^, could prove therapeutic to cocaine-using individuals, even individuals in the advanced stages of CUD.

Here, we employed locomotor behavioral assays to directly interrogate the cocaine-elicited behavioral adaptations to our MSN-targeted delivery of ZFP189 TFs, as well as operant cocaine IVSA of mice receiving ZFP189 TFs intra-NAc. Additionally, in other published work, we have also utilized reward-associated behavioral assays including conditioned place preference (CPP) and IVSA for cocaine in conjunction with NAc ZFP189 manipulations. By utilizing CRISPR-mediated activation of *Zfp189* expression in whole NAc, we observe that mice elect to self-administer more infusions of cocaine via IVSA and form less robust reward-associations for cocaine via CPP, and these altered conditioned behaviors were determined to result from *Zfp189* expression in *Drd2+* MSNs^1^. In IVSA experiments shown here where we utilized ZFP189 TFs delivered to whole NAc, we discovered that ZFP189^WT^ facilitates an increase of cocaine IVSA in a fixed ratio schedule of operant responding after a period of forced abstinence, whereas ZFP189^VPR^ impedes any increase in cocaine IVSA behaviors (Fig. 5). These operant and conditioned behavioral data further reinforce the notion that normal KZFP function, achieved by CRISPR-mediated activation of *Zfp189* or viral delivery of ZFP189^WT^ to the NAc, enables the brain changes necessary to facilitate the progression of CUD-related behaviors, including increases in cocaine taking. Notably, impeding normal KZFP function, achieved by virally delivering ZFP189^VPR^ to the NAc, halts these increases in cocaine taking behaviors, just as it halts cocaine-induced transcription seen in the snRNAseq (Fig. 4). Thus, ZFP189^VPR^ may prove a useful research tool to illuminate molecular strategies to slow, or even reverse, the brain transcriptional drivers of CUD.

We possess an incomplete picture for the natural biological mechanisms that control cocaine-induced TE release and transposition, which we discover disproportionately occurs in NAc MSNs immediately following the first cocaine exposure (Fig. 1). It is possible that some aspect of cocaine experience removes the constitutive epigenetic and transcriptional repression of TEs^44–50^ which might explain the observed TE expression. Alternatively, there may be unknown molecular mechanisms that actually activate TE expression, which could help explain the rapid temporal dynamics of TE release. Exploring the molecular origin of this phenomenon is an important future direction. Also, the cell-type specific molecular consequences of TE release and transposition, including genomic insertion and the lasting consequences on gene expression and genome integrity within *Drd1+* and *Drd2+* MSNs, will add additional dimensions to how KZFPs and TEs participate in the molecular pathogenesis of CUD. This is important to pursue, as genomic insertions as a result of TE transposition have been identified in the brains of individuals with CUD^36^, but the functional consequences of TE transposition, especially at the single-cell level, have not been well studied.

Lastly, we observe that ZFP189^WT^ functions similarly in both *Drd1+* and *Drd2+* MSNs to enhance the cocaine-induced expression of immune-related transcripts, while ZFP189^VPR^ impedes the expression of these same gene profiles (Fig. 4). This is consistent with the discovery that TEs have evolved to become *cis-*regulatory genomic domains primarily for immune genes^51,52^, as well as being able to influence neuroimmune response more generally^53–55^. In our other work virally delivering these ZFP189 TFs to the prefrontal cortex, we observed a similar impact on immune transcripts, yet were unable to determine the cell-type in which these transcripts were expressed^2^. Here, we discover that KZFP function directly controls gene expression entirely within NAc MSNs, which is summarized in the graphic in Figure 4e. These data point to KZFP-facilitated expression of immune genes in NAc MSNs as a molecular language for encoding response to cocaine experience. This is supported by considerable and growing evidence describing that cocaine use modulates immune signaling in brain reward regions^56–63^ and alters the neuroplasticity central to SUDs^64,65^. Thus, TEs may play an under-appreciated role in facilitating brain immune response to salient experience, like substance use.

Taken together, our research builds on the distinct observations that substance use is associated with brain expression of TEs^34,35,66^, KZFPs^1,67^, and immune factors^57,62,68,69^, and unifies these observations into a coherent and causative mechanism where the KZFP-mediated stabilization of TEs in brain MSNs enables cocaine-induced immune gene expression profiles and subsequent lasting molecular and behavioral adaptations to substance use experience. While growing evidence points to the co-option of genomic TEs as enabling transcriptional complexity and driving adaptations that have enabled evolutionary advantages to the host organism^70–74^, here we outline a mechanism by which the enhanced transcriptional diversity, facilitated by KZFP-mediated control of genomic TEs within NAc MSNs, causally drives cocaine-induced brain adaptations and may contribute to CUD pathogenesis. This adds nuance to discussions on how TEs contribute to higher-order cognitive functions and how aberrant brain KZFP function may confer risk for psychiatric syndromes, including SUDs.

## Methods

### 1. Animals

Male and female C57BL/6J mice, aged 8 to 12 weeks old, were purchased from Jackson Laboratories, and *Drd1*-Cre and *Drd2*-Cre bacterial artificial chromosome mice^75^ were bred in-house at the VCU transgenic mouse core. Mice were group housed (5 mice/cage) on a 12-hour light/dark cycle (lights on at 6am/off at 6pm) with food and water freely available. All tests were conducted in the light cycle. All mice were used in accordance with protocols approved by the Institutional Care and Use Committees at Virginia Commonwealth University School of Medicine.

### 2. Viral Packaging

We *de novo* synthesized ZFP189^NFD^, ZFP189^VPR^, and ZFP189^WT^ and sub-closed into either P1005 or a P1005 LS1L herpes simplex virus expression plasmids via ThermoFisher Scientific gateway LR Clonase II cloning reaction and Gateway LR Clonase II Enzyme mix kit (Cat # 11791-020 and 11971-100). Colonies were Maxiprepped (Qiagen Cat # 12163) and shipped to the Gene Delivery Technology Core at Massachusetts General Hospital for HSV packaging. Once packaged, aliquots were made and stored in −80° C to be used in viral gene transfer through stereotaxic surgery.

### 3. Stereotaxic Infusions

Stereotaxic surgeries targeting the NAc were performed as previously described^1,2,76^. Mice were anesthetized with I.P. injection of ketamine (100mg/kg) and xylazine (10mg/kg) dissolved in a sterile saline solution. Mice were then placed in a small animal stereotaxic device (Kopf Instruments) and the skull surface was exposed. 33-gauge needles (Hamilton Syringe) were utilized to infuse 1.0μL of virus at a rate of 0.2μL/minute followed by a 5-minute rest period to prevent backflow. The following coordinates were used to target the NAc: Bregma: anterior-posterior: +1.62mm, medial-lateral: 1.5mm, dorsal-ventral −4.4mm, 10° angle^1,76,77^. These coordinates combined with the 1.0 μL of virus transduces the core and shell of the NAc. All procedures occurred in our biosafety level 2+ facility.

### 4. Behavioral Paradigms

#### a. Locomotion Assay

Mice were either given an I.P. injection of cocaine (10mg/kg) or I.P. injection of vehicle saline and immediately placed in open field boxes for a locomotor sensitization trial of 30 minutes to freely move around the chamber, with distance recorded. Locomotor activity experiment sessions were tracked through EthoVision tracking software for analysis, captured as total distance moved in centimeters. Mice were placed in the same locomotor boxes for the duration of the experiment. Trials took place once per day at mid-day for seven days.

#### b. Cumulative Dose Response

The same open field boxes used in the locomotor activity paradigms were used for the locomotor dose response. Mice were allowed to habituate to their individual boxes for a 30-minute time period. Mice were then given an I.P. injection of saline and subjected to a 30-minute locomotor activity trial. This same timeline was used with subsequent doses of 1, 2.2, 6.8, and 25.2mg/kg cocaine injections, compounding on each other for total cumulative doses of 1, 3.2, 10, and 32 mg/kg I.P. injections of cocaine, all immediately followed by a 30-minute locomotor activity trial. Mice were placed in the same locomotor boxes for each trial, and trials took place at mid-day. Distance was measured (cm) and total distance within each ZFP189 variant group (ZFP189^WT^ or ZFP189^VPR^) were normalized against the ZFP189^NFD^ control and shown as a percent change.

### 5. Microscopy

Mice were transcardially perfused with 0.1M sodium phosphate buffer, followed by 4% paraformaldehyde (PFA) in 0.1M phosphate buffer. Brains were removed and post-fixed in 4% PFA overnight at 4°C. Following post-fix, brains were stored in 15% sucrose in phosphate buffer with 0.05% sodium azide for 24 hours, and then stored in 30% sucrose in phosphate buffer with 0.05% sodium azide until sectioning. Coronal sections at 40μm thick containing the NAc were then prepared on a cryostat (Leica CM1860). Sections were then mounted and cover slipped using ProLong Gold Antifade. Slides were kept at 4°C until imaging in a light-blocking slide box. Sections were imaged and captured with a Keyence BZX-800 fluorescent microscope. Slides were imaged with a 63x oil-immersion magnification.

### 6. Dendritic Spine Analysis

Images were captured on a Keyence BZX-800 fluorescent microscope, and individual dendritic segments were focused on and scanned at 0.7μm intervals along the z axis to obtain a z-stack. After capture, all images were performed in a two-dimensional projection using imageJ (NIH). 20μm in length of dendrite were analyzed on captured neurons. For each group, 2-6 20μm segments were analyzed from 4-5 mice. We operationally divided spines into three categories; (1) mushroom-like spines were dendritic protrusions with a head diameter > 0.5μm or > 2x the spine neck diameter; (2) stubby spines were dendritic protrusions with no discernable head and a length of ≤ 0.5μm; and (3) thin/filopodia-like spines were dendritic protrusions with a length of > 0.5μm and head diameter < 0.5μm or no discernable head^2,43^.

### 8. Tissue Collection, Nuclei Isolation from Mouse NAc

Mice were virally delivered HSV-ZFP189^NFD^, -ZFP189^WT^, -ZFP189^VPR^ intra-NAc to be used in snRNAseq analysis. Mice were given an I.P. injection of cocaine (10mg/kg) and immediately placed in open field boxes for a locomotor sensitization trial of 30 minutes to freely move around the chamber. Trials took place once per day at mid-day for 7 days. On the 7th day, one hour after I.P. injection of cocaine (10mg/kg), mice were euthanized by cervical dislocation without anesthesia and then were rapidly decapitated for brain removal. The brain was rapidly removed and blocked into 1mm-thick coronal sections using ice-cold brain matrices (Zivic instruments, Pittsburgh PA). Bilateral tissue punches from the nucleus accumbens were collected (NAc; 2 x14 gauge, internal diameter 1.6mm, n = 15 per sex, per ZFP189 variant). Tissue from each variant group was pooled together into one collection tube and rapidly frozen on dry ice. Tissue was then stored at −80°C until the day of nuclei isolation and sequencing. Nuclei were then isolated following the 10X Genomics protocol on nuclei isolation of adult mouse brain tissue for single nuclei RNA sequencing. Briefly, tissue lysis and nuclei washing, myelin removal, and density gradient centrifugation were performed for nuclei isolation. Once nuclei were successfully isolated, we followed the 10x Genomics Chromium single cell 3’ gene expression pipeline. The nuclei suspension was loaded into a Chromium Chip B with partitioning oil, reverse transcription reagents, and gel beads containing 10X Genomics barcodes. Barcoded cDNA was generated for each sample then pooled together. Library construction and sequencing on an Illumina HiSeq series sequencer was performed at a depth of approximately 50,000 reads per nuclei.

### 9. snRNAseq and analysis

#### a. Custom Reference Genome Generation

To incorporate three different viral constructs into the reference genome for single-nucleus RNA sequencing analysis, we generated custom reference genomes for each viral construct (ZFP189^VPR^, ZFP189^WT^, and ZFP189^NFD^), following steps recommended by 10X Genomics. In brief, for each viral construct the viral construct sequence was added to the mouse reference genome (mm39) to create a combined fasta file. A corresponding gene annotation in GTF format was also created. The length of each viral sequence was calculated, and an exon annotation was added to the existing GENCODE vM32 annotation file to produce a comprehensive annotation file.

#### b. Data Preprocessing and Quality Control

The sequencing data were processed using Cell Ranger software (version 7.1.0). The FASTQ files were aligned to the custom mouse reference genomes to generate feature-barcode matrices, and were further analyzed using the Seurat package (version 4.3). Initial quality control was performed to filter out low-quality nuclei (etc. number of features > 700, number of read > 3162, ratio of log10 features per UMI > 0.8, and percentage of mitochondrial gene expression < 5%).

#### c. Dimensionality Reduction and Clustering

Filtered data were normalized and log-transformed. Principal Component Analysis (PCA) was performed to reduce dimensionality, and the first 50 principal components were selected for downstream analysis. The selected components were used for Uniform Manifold Approximation and Projection (UMAP) to visualize the data in two dimensions. Graph-based clustering algorithms were applied to identify distinct cell-type populations. The known NAc cell-type clusters were annotated using canonical gene markers in rodents^21,78,79^.

#### d. Differential Expression Analysis

Differential expression was compared between ZFP189 variants (ZFP189^VPR^ or ZFP189^WT^) and the control (ZFP189^NFD^) using the general linear model provided in MAST package^80^. Genes and TEs were tested for differential expression when expressed in minimal 10% of the nuclei in at least one group, with a minimum difference of 5% in the percentage of expressing cells between the two groups. Genes with an FDR adjusted *P*-value < 0.05 and an absolute Log_2_(fold change) > 0.25 were considered significantly differentially expressed genes (DEGs).

#### e. Ingenuity Pathway Analysis

To assess the function of the significant DEGs regulated by ZFP189 TFs, we performed an ingenuity pathway analysis (IPA)^81^. The networks were generated through the use of Qiagen IPA (QIAGEN Inc., https://digitalinsights.qiagen.com/IPA). Significant DEGs from *Drd1*+ and *Drd2*+ MSNs by ZFP189 variant were analyzed using IPA to determine associated pathways. Significantly regulated pathways (FDR< 0.05) were sorted by z-score and graphed into a regulatory heatmap using SRplot matrix heat map generator (https://www.bioinformatics.com.cn/srplot).

#### f. Rank-rank hypergeometric overlap

RRHO allows for the direct comparison of gene expression profiles in a threshold-free capacity^82^. Threshold-free differential gene lists were used to generate RRHOs comparing *Drd1+* and *Drd2+* MSN cell types between ZFP189 TF variants (ZFP189^VPR^ or ZFP189^WT^). RRHO plots were generated in R using the RRHO package (https://bioconductor.org/packages/release/bioc/html/RRHO.html).

#### g. union heat map

Union heat maps were generated from a full list of differentially expressed genes (FDR adjusted *P-*value < 0.05 and an absolute Log_2_(fold change) > 0.25) identified within transfection of ZFP189 variants (ZFP189^VPR^ or ZFP189^WT^) within the *Drd1+* and *Drd2+* MSN cell populations. Log_2_(fold change) of DEGs were then graphed using Morpheus (https://software.broadinstitute.org/morpheus/).

#### h. Identification of Transposable Elements

To identify and analyze TEs in snRNA-seq data, we utilized the SoloTE tool (version 1.06). The RepeatMasker output for the mouse genome (mm39) was firstly converted to TE annotations for SoloTE. The position-sorted read alignment generated by Cell Ranger in an aforementioned step was quantified with the TE annotations and the default parameters of SoloTE.

#### i. TE and DEG Analysis of Previously Published SnRNAseq Datasets

Publicly available snRNAseq datasets were downloaded from NCBI GEO (<u>GSE137763</u>, <u>GSE222418</u>). Similar to previous analyses described above, TEs were identified using the SoloTE tool (version 1.06). The RepeatMasker output for the rat genome (Ensembl Rnot_6.0 for the acute cocaine dataset^21^ and Ensembl mRatBn7.2 for the repeated cocaine dataset^39^ based on different genome updates when papers were published) was converted to TE annotations for SoloTE. The position-sorted read alignment generated by Cell Ranger in an aforementioned step was quantified with the TE annotations and the default parameters of SoloTE. Similarly, DEGs were generated against the saline treated control rats, using the general linear model provided in MAST package^80^. Genes and TEs were tested for differential expression when expressed in minimal 10% of the nuclei in at least one group, with a minimum difference of 5% in the percentage of expressing cells between the two groups. Genes with an FDR adjusted *P*-value < 0.05 and an absolute Log_2_(fold change) > 0.25 were considered DEGs and DETEs.

### 10. Cocaine self-administration procedures

#### a. Jugular vein catheterization

Mice were prepared for surgery by removing fur at the surgical site with electrical clippers. The skin was then aseptically prepared with alternating applications of 70% ethanol and chlorohexidine solution. An incision was made over the right jugular vein, until the vein was exposed. Once the vein was exposed, a small incision was made in the vein for the catheter to be inserted into, then secured by a non-absorbable suture. Next, an incision was made in the intrascapular region, for the free end of the catheter to be subcutaneously tunneled and exteriorized. The catheters were then secured to the vascular access button (VAB) and then filled with a heparin/glycerol lock solution. Finally, a silicone mesh disk portion of the VAB is placed subcutaneously. Local anesthetics were applied topically to the incision sites and closed with non-absorbable sutures.

### b. Jugular vein catheter maintenance

Catheters were maintained through a routine of sterilization that involved wiping the VAB with ethanol-soaked cotton swabs before and garter being placed in the IVSA chambers, and flushing daily with a 0.05 ml heparinized saline solution (30 U) containing ampicillin antibiotic (0.5 mg/ml) via a pin port tipped syringe. During periods of forced abstinence, a glycerin-based locking solution was used to seal the catheters, with 0.01ml being pumped into the VAB. Mice could remain locked for a maximum of 7 days before requiring being unlocked and flushed with the heparinized solution described above. To unlock the VAB, this locking solution was drawn up via an empty pin port tipped syringe until a small amount of blood was drawn to indicate patency.

#### c. Anesthetization and surgery of catheterized mice

Intra-NAc infusion of ZFP189 TFs via stereotaxic surgery was consistent with the procedure described above in “Stereotaxic Infusions” section, with one additional step. To avoid damaging the jugular vein catheters, mice were briefly anesthetized (5-10 seconds) using a bell jar with isoflurane to allow for the I.P. injection of ketamine and xylazine solution without needing to scruff the animals. The rest of the surgical procedure continues as described above.

#### d. Operant chambers

The operant chambers used in this study were from med-associates, classic modular test chambers (ENV-307A-CT), placed in standard MDF sound attenuating cubicles (ENV-022MD). The operant chambers contained a LED house light (ENV-315M-LED) that would be on during testing sessions, two retractable levers (ENV-312-3M) along with two cue induced lights to indicate active and inactive levers (ENV-321M). The operant chamber was also paired with a pellet dispenser (ENV-203-20) and pellet receptacle (ENV-303M) which allowed for the administration of sucrose pellets in food training. This setup also included a variable speed pump (PHM-210) that allowed for the infusion of cocaine through tubing to the catheters implanted in the mice.

#### e. Cocaine infusions via syringe pump

Each infusion of cocaine was 0.5mg/kg/infusion. To achieve this, syringes were filled with 0.25 mg/ml of cocaine in saline, which would be delivered with the med-associates syringe pump running at a rate of 3.33 RPM for 2.82 seconds. This rate was calculated based on the manufacturer’s recommendation. This dosage was consistent throughout all cocaine IVSA experiments across all sessions.

#### f. Training-Cocaine IVSA fixed ratio 1 scheduled study

Mice were trained on cocaine training sessions lasting three hours on a FR1 reinforcement schedule. The active lever was indicated via a stim-light illuminated above, which would turn off during the timeout period. Active lever presses during timeouts were recorded, but did not result in infusions. Similarly inactive lever presses throughout the entire session did not result in an infusion. During the first two sessions, infusions were capped at a maximum of 60, with a timeout period of 15 seconds following each infusion to avoid potential overdose. At the start of the third day, the cap on infusions was elevated to 120 per session, but the timeout period remained unchanged. Mice remained in this training until they showed an ability to discriminate between active and inactive levers and a stable response was achieved, defined as individual animal infusions that did not differ by more than 35% from the average of the last two sessions, inclusive, or the group average did not vary by more than 20% over the last two sessions, inclusive. Generally, this process took between 7 and 10 days. Following the achievement of stable responding rates, mice would undergo the intra-NAc infusion of ZFP189 TFs.

#### g. Testing-Cocaine IVSA fixed ratio 1 study

All testing sessions were three hours in duration and remained on a FR1 reinforcement schedule. Following surgery testing consisted of 14-consecutive sessions. At the end of the 14 sessions, mice had their catheters locked with the method described above in the “Jugular vein catheter maintenance” section and underwent seven days of forced abstinence without access to cocaine. Following the seven days of forced abstinence mice returned to the chambers for seven more consecutive sessions.

## Contributions

GMS and PJH conceived and conceptualized experimental design and data interpretation; GMS, JAP, ALK, CRP, DPL, RKK, NLT, XQ, and PJH performed all experimentation; GMS and PJH analyzed all RNA sequencing data; RLN packaged herpes simplex viruses; GMS and PJH contributed to manuscript preparation; and all authors reviewed and approved final manuscript draft.

## Corresponding author

Correspondence to Peter J. Hamilton

## Acknowledgements

This work was supported by the NIH grants R00DA045795, P30DA033934, R34DA061267, R01DA058958, R01DA058089, and Blink Scholar funds to PJH, as well as T32DA007027 and T32GM148403 to GMS. Microscopy was performed at the VCU Microscopy Facility, supported, in part, by funding from the NIH-NCI Cancer Center Support Grant P30CA016059. Transgenic mouse lines used in this project were provided by the VCU Massey Comprehensive Cancer Center Transgenic/Knockout Mouse Shared Resource, supported, in part, with funding from the NIH-NCI Cancer Center Support Grant P30CA16059.

## Data Availability

Raw and processed snRNAseq and RNAseq gene expression data will be made available via Gene Expression Omnibus data upon publication. Other data that supports the findings from this study are available from the corresponding author upon request.

## References

1. Teague, C. D. et al. CREB binding at the Zfp189 promoter within medium spiny neuron subtypes differentially regulates behavioral and physiological adaptations over the course of cocaine use. Biol Psychiatry 93, 502–511 (2023).

2. Truby, N. L. et al. A zinc finger transcription factor enables social behaviors while controlling transposable elements and immune response in prefrontal cortex. Transl Psychiatry 14, 59 (2024).

3. de Tribolet-Hardy, J. et al. Genetic features and genomic targets of human KRAB-zinc finger proteins. Genome Res 33, 1409–1423 (2023).

4. Wolf, G. et al. KRAB-zinc finger protein gene expansion in response to active retrotransposons in the murine lineage. eLife 9, e56337 (2020).

5. Playfoot, C. J. et al. Transposable elements and their KZFP controllers are drivers of transcriptional innovation in the developing human brain. Genome Res 31, 1531–1545 (2021).

6. Pontis, J. et al. Hominoid-Specific Transposable Elements and KZFPs Facilitate Human Embryonic Genome Activation and Control Transcription in Naive Human ESCs. Cell Stem Cell 24, 724–735.e5 (2019).

7. Abuse, N. I. on D. Drug Overdose Death Rates | National Institute on Drug Abuse (NIDA). https://nida.nih.gov/research-topics/trends-statistics/overdose-death-rates (2024).

8. Day, J. J., Roitman, M. F., Wightman, R. M. & Carelli, R. M. Associative learning mediates dynamic shifts in dopamine signaling in the nucleus accumbens. Nat Neurosci 10, 1020– 1028 (2007).

9. Schultz, W., Apicella, P. & Ljungberg, T. Responses of monkey dopamine neurons to reward and conditioned stimuli during successive steps of learning a delayed response task. J. Neurosci. 13, 900–913 (1993).

10. Schultz, W. Dopamine reward prediction error coding. Dialogues in Clinical Neuroscience 18, 23–32 (2016).

11. Nestler, E. J. The Neurobiology of Cocaine Addiction. Sci Pract Perspect 3, 4–10 (2005).

12. Dong, Y. et al. CREB modulates excitability of nucleus accumbens neurons. Nat Neurosci 9, 475–477 (2006).

13. Renthal, W. et al. Genome Wide Analysis of Chromatin Regulation by Cocaine Reveals a Novel Role for Sirtuins. Neuron 62, 335–348 (2009).

14. Lobo, M. K. et al. Cell type-specific loss of BDNF signaling mimics optogenetic control of cocaine reward. Science 330, 385–390 (2010).

15. Hikida, T., Kimura, K., Wada, N., Funabiki, K. & Nakanishi, S. Distinct roles of synaptic transmission in direct and indirect striatal pathways to reward and aversive behavior. Neuron 66, 896–907 (2010).

16. Tai, L.-H., Lee, A. M., Benavidez, N., Bonci, A. & Wilbrecht, L. Transient stimulation of distinct subpopulations of striatal neurons mimics changes in action value. Nat Neurosci 15, 1281–1289 (2012).

17. Calipari, E. S. et al. In vivo imaging identifies temporal signature of D1 and D2 medium spiny neurons in cocaine reward. Proc Natl Acad Sci U S A 113, 2726–2731 (2016).

18. Heinsbroek, J. A. et al. Loss of Plasticity in the D2-Accumbens Pallidal Pathway Promotes Cocaine Seeking. J Neurosci 37, 757–767 (2017).

19. van Zessen, R. et al. Dynamic dichotomy of accumbal population activity underlies cocaine sensitization. eLife 10, e66048 (2021).

20. Hughes, B. W. et al. NPAS4 supports cocaine-conditioned cues in rodents by controlling the cell type-specific activation balance in the nucleus accumbens. Nat Commun 15, 5971 (2024).

21. Savell, K. E. et al. A dopamine-induced gene expression signature regulates neuronal function and cocaine response. Sci Adv 6, eaba4221 (2020).

22. Levin, H. L. & Moran, J. V. Dynamic interactions between transposable elements and their hosts. Nat Rev Genet 12, 615–627 (2011).

23. Bourque, G. et al. Ten things you should know about transposable elements. Genome Biol 19, 199 (2018).

24. Imbeault, M., Helleboid, P.-Y. & Trono, D. KRAB zinc-finger proteins contribute to the evolution of gene regulatory networks. Nature 543, 550–554 (2017).

25. Helleboid, P. et al. The interactome of KRAB zinc finger proteins reveals the evolutionary history of their functional diversification. EMBO J 38, e101220 (2019).

26. Takahashi, T. et al. LINE-1 activation in the cerebellum drives ataxia. Neuron 110, 3278–3287.e8 (2022).

27. Chen, Y.-C., Maupas, A. & Nowick, K. Regulatory networks of KRAB zinc finger genes and transposable elements changed during human brain evolution and disease. 2023.12.18.569574 Preprint at 10.1101/2023.12.18.569574 (2024).

28. Wolf, D. & Goff, S. P. Embryonic stem cells use ZFP809 to silence retroviral DNAs. Nature 458, 1201–1204 (2009).

29. Jacobs, F. M. et al. An evolutionary arms race between KRAB zinc finger genes 91/93 and SVA/L1 retrotransposons. Nature 516, 242–245 (2014).

30. Najafabadi, H. S., Albu, M. & Hughes, T. R. Identification of C2H2-ZF binding preferences from ChIP-seq data using RCADE. Bioinformatics 31, 2879–2881 (2015).

31. Schmitges, F. W. et al. Multiparameter functional diversity of human C2H2 zinc finger proteins. Genome Res 26, 1742–1752 (2016).

32. Bruno, M., Mahgoub, M. & Macfarlan, T. S. The Arms Race Between KRAB–Zinc Finger Proteins and Endogenous Retroelements and Its Impact on Mammals. Annu. Rev. Genet. 53, 393–416 (2019).

33. Sundaram, V. & Wysocka, J. Transposable elements as a potent source of diverse cis-regulatory sequences in mammalian genomes. Philos Trans R Soc Lond B Biol Sci 375, 20190347 (2020).

34. Okudaira, N., Ishizaka, Y. & Nishio, H. Retrotransposition of Long Interspersed Element 1 Induced by Methamphetamine or Cocaine. J Biol Chem 289, 25476–25485 (2014).

35. Maze, I. et al. Cocaine dynamically regulates heterochromatin and repetitive element unsilencing in nucleus accumbens. Proc Natl Acad Sci U S A 108, 3035–3040 (2011).

36. Doyle, G. A. et al. Reading LINEs within the cocaine addicted brain. Brain Behav 7, e00678 (2017).

37. Moszczynska, A., Flack, A., Qiu, P., Muotri, A. R. & Killinger, B. A. Neurotoxic Methamphetamine Doses Increase LINE-1 Expression in the Neurogenic Zones of the Adult Rat Brain. Sci Rep 5, 14356 (2015).

38. Rodríguez-Quiroz, R. & Valdebenito-Maturana, B. SoloTE for improved analysis of transposable elements in single-cell RNA-Seq data using locus-specific expression. Commun Biol 5, 1063 (2022).

39. Phillips, R. A. et al. Distinct subpopulations of D1 medium spiny neurons exhibit unique transcriptional responsiveness to cocaine. Mol Cell Neurosci 125, 103849 (2023).

40. Hamilton, P. J. et al. Cell-Type-Specific Epigenetic Editing at the Fosb Gene Controls Susceptibility to Social Defeat Stress. Neuropsychopharmacol. 43, 272–284 (2018).

41. Lardner, C. K. et al. Gene-targeted, CREB-mediated induction of ΔFosB controls distinct downstream transcriptional patterns within D1 and D2 medium spiny neurons. Biol Psychiatry 90, 540–549 (2021).

42. Graziane, N. M. et al. Opposing mechanisms mediate morphine- and cocaine-induced generation of silent synapses. Nat Neurosci 19, 915–925 (2016).

43. Wright, W. J. et al. Silent Synapses Dictate Cocaine Memory Destabilization and Reconsolidation. Nat Neurosci 23, 32–46 (2020).

44. Grassi, D. A., Jönsson, M. E., Brattås, P. L. & Jakobsson, J. TRIM28 and the control of transposable elements in the brain. Brain Research 1705, 43–47 (2019).

45. Rowe, H. M. et al. KAP1 controls endogenous retroviruses in embryonic stem cells. Nature 463, 237–240 (2010).

46. Turelli, P. et al. Interplay of TRIM28 and DNA methylation in controlling human endogenous retroelements. Genome Res 24, 1260–1270 (2014).

47. Collins, P. L., Kyle, K. E., Egawa, T., Shinkai, Y. & Oltz, E. M. The histone methyltransferase SETDB1 represses endogenous and exogenous retroviruses in B lymphocytes. Proc Natl Acad Sci U S A 112, 8367–8372 (2015).

48. Ecco, G., Imbeault, M. & Trono, D. A tale of domestication: the endovirome, its polydactyl controllers and the species-specificity of human biology. Development 144, 2719–2729 (2017).

49. Rowe, H. M. et al. De novo DNA methylation of endogenous retroviruses is shaped by KRAB-ZFPs/KAP1 and ESET. Development 140, 519–529 (2013).

50. Elsässer, S. J., Noh, K.-M., Diaz, N., Allis, C. D. & Banaszynski, L. A. Histone H3.3 is required for endogenous retroviral element silencing in embryonic stem cells. Nature 522, 240–244 (2015).

51. Ye, M. et al. Specific subfamilies of transposable elements contribute to different domains of T lymphocyte enhancers. Proc Natl Acad Sci U S A 117, 7905–7916 (2020).

52. Chuong, E. B., Elde, N. C. & Feschotte, C. Regulatory evolution of innate immunity through co-option of endogenous retroviruses. Science 351, 1083–1087 (2016).

53. Ochoa, E. et al. Pathogenic tau–induced transposable element–derived dsRNA drives neuroinflammation. Sci Adv 9, eabq5423.

54. Saleh, A., Macia, A. & Muotri, A. R. Transposable Elements, Inflammation, and Neurological Disease. Front Neurol 10, 894 (2019).

55. Volkman, H. E. & Stetson, D. B. The enemy within: endogenous retroelements and autoimmune disease. Nat Immunol 15, 415–422 (2014).

56. Murakami, G. et al. MHC class I in dopaminergic neurons suppresses relapse to reward seeking. Sci Adv 4, eaap7388 (2018).

57. Brown, K. T. et al. Innate immune signaling in the ventral tegmental area contributes to drug-primed reinstatement of cocaine seeking. Brain Behav Immun 67, 130–138 (2018).

58. Northcutt, A. L. et al. DAT isn’t all that: cocaine reward and reinforcement requires Toll Like Receptor 4 signaling. Mol Psychiatry 20, 1525–1537 (2015).

59. Guo, M.-L. et al. Cocaine-mediated microglial activation involves the ER stress-autophagy axis. Autophagy 11, 995–1009 (2015).

60. Liao, K. et al. Cocaine-mediated induction of microglial activation involves the ER stress-TLR2 axis. J Neuroinflammation 13, 33 (2016).

61. Little, K. Y. et al. Decreased brain dopamine cell numbers in human cocaine users. Psychiatry Research 168, 173–180 (2009).

62. Sorg, B. A., Krueger, J. M., Churchill, L., Cearley, C. N. & Blindheim, K. Acute Cocaine Increases Interleukin-1β mRNA and Immunoreactive Cells in the Cortex and Nucleus Accumbens. Neurochem Res 36, 686–692 (2011).

63. Zhu, R. et al. Toll-like receptor 3 modulates the behavioral effects of cocaine in mice. J Neuroinflammation 15, 93 (2018).

64. Kashima, D. T. & Grueter, B. A. Toll-like receptor 4 deficiency alters nucleus accumbens synaptic physiology and drug reward behavior. Proc Natl Acad Sci U S A 114, 8865–8870 (2017).

65. Lewitus, G. M. et al. Microglial TNFα suppresses cocaine-induced plasticity and behavioral sensitization. Neuron 90, 483–491 (2016).

66. Moszczynska, A. Differential Responses of LINE-1 in the Dentate Gyrus, Striatum and Prefrontal Cortex to Chronic Neurotoxic Methamphetamine: A Study in Rat Brain. Genes (Basel) 11, 364 (2020).

67. Phan, B. N. et al. Single nuclei transcriptomics in human and non-human primate striatum in opioid use disorder. Nat Commun 15, 878 (2024).

68. Browne, C. J. et al. Transcriptional signatures of heroin intake and relapse throughout the brain reward circuitry in male mice. Science Advances 9, eadg8558 (2023).

69. Mews, P. et al. Convergent abnormalities in striatal gene networks in human cocaine use disorder and mouse cocaine administration models. Science Advances 9, eadd8946 (2023).

70. Xia, B. et al. On the genetic basis of tail-loss evolution in humans and apes. Nature 626, 1042–1048 (2024).

71. Laperriere, D., Wang, T.-T., White, J. H. & Mader, S. Widespread Alu repeat-driven expansion of consensus DR2 retinoic acid response elements during primate evolution. BMC Genomics 8, 23 (2007).

72. Kvikstad, E. M. & Makova, K. D. The (r)evolution of SINE versus LINE distributions in primate genomes: Sex chromosomes are important. Genome Res 20, 600–613 (2010).

73. Rowe, H. M. et al. TRIM28 repression of retrotransposon-based enhancers is necessary to preserve transcriptional dynamics in embryonic stem cells. Genome Res 23, 452–461 (2013).

74. Senft, A. D. & Macfarlan, T. S. Transposable elements shape the evolution of mammalian development. Nat Rev Genet 22, 691–711 (2021).

75. Gong, S. et al. Targeting Cre Recombinase to Specific Neuron Populations with Bacterial Artificial Chromosome Constructs. J Neurosci 27, 9817–9823 (2007).

76. Hamilton, P. J., Lim, C. J., Nestler, E. J. & Heller, E. A. Viral Expression of Epigenome Editing Tools in Rodent Brain Using Stereotaxic Surgery Techniques. Methods Mol Biol 1767, 205–214 (2018).

77. Hamilton, P. J., Lim, C. J., Nestler, E. J. & Heller, E. A. Stereotaxic Surgery and Viral Delivery of Zinc-Finger Epigenetic Editing Tools in Rodent Brain. Methods Mol Biol 1867, 229–238 (2018).

78. Chen, R. et al. Decoding molecular and cellular heterogeneity of mouse nucleus accumbens. Nat Neurosci 24, 1757–1771 (2021).

79. Gokce, O. et al. Cellular Taxonomy of the Mouse Striatum as Revealed by Single-Cell RNA-Seq. Cell Rep 16, 1126–1137 (2016).

80. Finak, G. et al. MAST: a flexible statistical framework for assessing transcriptional changes and characterizing heterogeneity in single-cell RNA sequencing data. Genome Biol 16, 278 (2015).

81. Krämer, A., Green, J., Pollard, J. & Tugendreich, S. Causal analysis approaches in Ingenuity Pathway Analysis. Bioinformatics 30, 523–530 (2014).

82. Plaisier, S. B., Taschereau, R., Wong, J. A. & Graeber, T. G. Rank–rank hypergeometric overlap: identification of statistically significant overlap between gene-expression signatures. Nucleic Acids Res 38, e169 (2010).

